# Hypercholesterolemia-induced impairment in sorafenib functionality is overcome by avasimibe co-treatment

**DOI:** 10.1101/2024.03.27.586757

**Authors:** Dipti Athavale, Himanshi Yaduvanshi, Firoz Khan Bhati, Shyamananda Singh Mayengbam, Tushar More, Srikanth Rapole, Manoj Kumar Bhat

**Affiliations:** National Centre for Cell Science, Department of Biotechnology, Government of India, Savitribai Phule Pune University Campus, Ganeshkhind, Pune-411 007, India

**Author notes:** Corresponding author: Dr Manoj Kumar Bhat, DBT Distinguished Scientist, National Centre for Cell Science, Pune, India. E-mail address (M.K. Bhat). Firoz Khan Bhati and Himanshi Yaduvanshi contributed equally to this article. **Email address of authors:** Dipti Athavale, Himanshi Yaduvanshi^#^, Firoz Khan Bhati^#^, Shyamananda Singh Mayengbam Tushar More, Srikanth Rapole Manoj Kumar Bhat.

**Keywords:** Hepatocellular carcinoma, Sorafenib, Hypercholesterolemia, Avasimibe

## Abstract

Avasimibe; a cholesterol-lowering drug with a proven safety in clinical trials, has recently been repositioned as an anticancer agent in various preclinical investigations. A study from our group reported that hypercholesterolemia promotes hepatocellular carcinoma (HCC) cell survival and hampers the anticancer effect of sorafenib, a kinase inhibitor. In the present study, we demonstrate that in HCC under hypercholesterolemic conditions the anticancer property of sorafenib is potentiated by avasimibe (AVA) co-treatment. Further, to elucidate the role of hypercholesterolemia on sorafenib efficacy, *in vitro* and *in vivo* models of HCC were used. *In vitro*, co-treatment of both drugs synergistically inhibited HCC cell viability and induced cell death under normal and hypercholesterolemic conditions. At the molecular level, downregulation of ERK signalling and induction of endoplasmic reticulum stress are likely to contribute to the combinatorial cytotoxic effect of sorafenib and avasimibe *in vitro*. In mice, fed on a high-cholesterol diet (HCD), the efficacy of sorafenib was restored by co-administration of AVA. Collectively, these findings suggest that impairment in the efficacy of sorafenib because of hypercholesterolemic phenotype could be restored by AVA co-treatment, which may have implications towards treatment strategy.

**Highlights:**

- Cholesterol impedes sorafenib efficacy in Hepatocellular carcinoma cells.
- Avasimibe restores the functionality of sorafenib under hypercholesterolemic environment.
- Combine treatment of sorafenib and avasimibe synergistically enhances cytotoxicity in hepatocellular carcinoma.
- Sorafenib and avasimibe treatment in the presence of LDLc.is associated with diminished ERK activation and increased ER stress.

## 1. Introduction

Hepatocellular carcinoma (HCC), a cancer originating from hepatocytes, accounts for more than 80% of all primary liver cancer cases. According to the International Agency for Research on Cancer (IARC), approximately 9,05,677 new cases of HCC were reported in 2020. HCC is the fifth most common cancer worldwide and is the third leading cause of cancer-related mortalities. According to the Globocan 2020 report, in India HCC ranked tenth among all cancers with 34,734 new cases reported. During the same period, 33,793 deaths were reported [1]. Cirrhosis development contributed by chronic hepatitis B or C virus infections and excess alcohol intake culminates into HCC development. Additionally, obesity and diabetes are arising as independent risk factors for non-alcoholic fatty liver disease, which also contributes to HCC development [3,4]. The selection of therapeutic interventions for HCC treatment depends on the stage of the disease. For small, localized tumors surgical resection or liver transplantation is recommended. For advanced and inoperable HCC with severely damaged liver functions; systemic targeted therapies or non-invasive therapies like transarterial chemoembolisation are advised [2].

Sorafenib (SORA) is approved as a front-line therapeutic agent for the treatment of advanced HCC. It is an orally administered tyrosine kinase inhibitor that blocks RAS/RAF/MEK/ERK kinases which are involved in proliferative and angiogenic pathways. As reported by a phase III clinical trial study, SORA increased the overall survival of HCC patients as compared to the placebo group. However; the survival period was unsatisfactory (∼3 months). Moreover, mechanisms of primary and acquired drug resistance are emerging which limits its clinical use. This raises the urgent need to develop effective therapeutic strategies for HCC, especially by combining SORA with other potential anticancer agents [5].

Elevated serum cholesterol levels (hypercholesterolemia) and altered intracellular cholesterol metabolism not only influence tumor progression but also hamper therapeutic outcomes in certain cancer types including HCC [6–10]. Commonly prescribed anti-hypercholesterolemic drugs like statins alone or in combination with other chemotherapeutic drugs have been shown to exhibit anti-proliferative and pro-apoptotic effects in various cancer types [11–13]. Cells esterify excess cholesterol with fatty acids forming cholesteryl esters (CE) and these are stored as lipid droplets (LD). This reaction is catalysed by an enzyme known as acyl-coenzyme A cholesterol acyl transferase (ACAT) or sterol O-acyltransferase. Avasimibe (AVA) is an inhibitor of ACAT and because of its cholesterol-lowering effects, it was being used for treating atherosclerosis [14–17].

Recently, AVA has been resuscitated with its newly explored anticancer effects in preclinical models. It is reported to inhibit ACAT-mediated aberrant accumulation of CE thereby reducing pancreatic tumor growth and its metastasis [18]. AVA in combination with gemcitabine synergistically exerts cytotoxic effects on drug-resistant pancreatic adenocarcinoma cells [19]. In the mouse melanoma model, it was reported that AVA potentiated the antitumor activity of CD8^+^ T cells to control the growth and metastasis of tumor [20].

By using appropriate *in-vitro* and *in-vivo* models in this study, the combinatorial effect of SORA and AVA was investigated in HCC. The combination of SORA and AVA synergistically inhibited survival and induced death in HCC cells in the presence of LDLc. Mechanistically, reduced ERK signalling and elevated ER stress were associated with their combinatorial effect. Collectively, these findings suggest that impairment in SORA efficacy because of hypercholesterolemic phenotype could be restored by AVA co-treatment which may have implications towards treatment strategies.

## 2. Materials and methods

### 2.1 Cell culture and reagents

Human hepatocellular carcinoma cell lines Hep-G2, Huh-7 and Hep3B were routinely cultured in Dulbecco’s modified Eagle medium (DMEM) with 10% heat-inactivated fetal bovine serum (Life Technologies, CA, USA). The culture medium was supplemented with penicillin (100 U/ ml), and streptomycin (100 μg/ml) (Invitrogen Life Technologies, CA, USA). Cells were maintained in a humidified atmosphere of 5% CO_2_ at 37 °C. Hep-G2 and Hep3B were procured from ATCC (Manassas, VA, USA) and were maintained at our in-house cell repository at the National Centre for Cell Science (NCCS), Pune, India. Huh-7 cell line was a generous gift from Dr. Ralf Bartenschlager (University of Heidelberg, Germany). Human low-density lipoprotein cholesterol (hLDLc) and high-density lipoprotein cholesterol (hHDLc) were purchased from Lee Biosolutions (Missouri, USA). Simvastatin (SIM) was procured from Enzo Life Sciences Inc (NY, USA). Avasimibe (AVA) and propidium iodide were purchased from Sigma-Aldrich (MO, USA). Primary antibodies; p-eIF2α (3398) and eIF2α (9722) were purchased from Cell Signalling Technology (MA, USA). Sorafenib (SORA) (sc-357801A), antibodies against PCNA (sc-56), β-actin (sc-1615), p-ERK (sc-7383), ERK (sc-154), mt-TFAM (sc-376672), and HRP-conjugated secondary antibodies were purchased from Santa Cruz Biotechnology (CA, USA).

### 2.2 MTT assay

Cells (2 × 10^3^) were seeded into 96 well plates and allowed to adhere for 24 h. These were treated with indicated concentrations of SORA, SIM and AVA for 48 h in DMEM with 5% FBS. To check the effect of hypercholesterolemia, cells were pre-incubated in DMEM (5% FBS) containing indicated concentrations of hLDLc or hHDLc, for 24 h followed by drug treatment as indicated in figure legends for 48 h. At the end of treatments, the medium was replaced with 50 µl of MTT solution (1 mg/ml) and cells were incubated for 4 h. Formazan crystals formed were dissolved by adding isopropanol (100 µl) and absorbance was measured in a microplate reader (MultiSkan Go, Thermo Fisher Scientific, OH, USA) at 570 nm. The absorbance of vehicle-treated cells was considered as 100 % cell survival. The efficacy of drug combination was evaluated by calculating the coefficient of drug interaction (CDI) as follows: CDI=AB/ (A/B); where AB is the ratio of the combination group to the control group. A and B are the ratio of the single agent group to the control group.

### 2.3 Long-term survival assay

Cells (5 × 10^2^) were seeded in 48-well plates and allowed to adhere. These were cultured with LDLc and treated with drugs as mentioned in respective figure legends. At the end of the treatment periods, the medium was replaced with fresh DMEM without drugs and cells were allowed to grow for a further 10 days by changing the medium after every second day. Subsequently, cells were processed for crystal violet staining as described before [10].

### 2.4 Propidium iodide (PI) staining

HCC cells (3 × 10^5^) were treated with drugs as mentioned in respective figure legends. At the end of the treatment period, cells were trypsinized and washed twice with ice-cold PBS by centrifugation at 1500 rpm at 4°C for 5 min. The supernatant was aspirated out and cells were resuspended in cold PBS containing PI (2 µg/ml) for 2 min followed by washing with cold PBS at 1500 rpm at 4°C for 5 min. PI staining was measured in the FL2 channel by using FACS Calibur (BD Bioscience, NJ, USA) and data were analyzed with CellQuest Pro software (BD Bioscience, NJ, USA).

### 2.5 LDH assay

HCC cells (3 × 10^5^) were plated in 35 mm dishes and allowed to adhere for 24 h. The cells were treated as described earlier. The supernatant medium was collected and LDH release was measured by LDH activity assay kit by Spinreact (Girona, Spain) as per the manufacturer’s instructions.

### 2.6 Mito Tracker Deep Red FM staining

Cells (1 × 10^3^) were seeded on coverslips in a 24-well plate and allowed to adhere for 24 h. Following the treatment as per the experimental requirement, cells were washed with DMEM without FBS. Mito Tracker™ Deep Red FM stock solution was diluted in DMEM without FBS to a final concentration of 100 nM and added to the wells. Cells were incubated at 37°C for 30 min. These were then washed with PBS and fixed. Subsequently, coverslips were mounted on slides with a mounting medium containing DAPI (Santa Cruz Biotechnology, CA, USA) and images were captured on a confocal microscope (Carl Zeiss, Heidelberg, Germany). Images were subsequently processed by LSM image analysis software (Carl Zeiss, Heidelberg, Germany).

### 2.7 Nile red staining

Cells (1 × 10^3^) were seeded on coverslips in a 24-well plate, allowed to adhere for 24 h and treated as mentioned in respective figure legends. At the end of the treatment period, cells were washed with PBS and fixed with 4% paraformaldehyde at RT for 10 min. Then cells were again washed with PBS. These fixed cells were stained with Nile Red (1 μg/ ml in PBS) at 37°C for 10 min followed by washing with PBS. Subsequently, coverslips were mounted on slides with a mounting medium containing DAPI (Santa Cruz Biotechnology, CA, USA). The images were captured on a confocal microscope and processed by LSM image analysis software (Carl Zeiss, Heidelberg, Germany).

### 2.8 Western blotting

Whole-cell lysates were resolved on SDS-PAGE and Western blotting was performed as described previously [10].

### 2.9 Intracellular SORA quantification

Intracellular SORA quantification was performed by liquid chromatography-multiple reactions monitoring mass spectrometry (LC-MRM/MS). Hep-G2 and Huh-7 cells (5 × 10^5^) were plated in 60 mm plates and pre-treated with LDLc (100 µg/ml) for 3 h. Thereafter, these were treated with SORA (5 µM) in the presence or absence of LDLc for 3 h. At the end of the treatment period, cells were washed with 1X PBS. To extract SORA, cells were homogenized in ice-cold methanol (400 µl) containing internal standard (L-phenylalanine) using zirconium beads. Homogenization was carried out using Precellys Homogenizer (Bertin Corp. MD, USA). The homogenization program was set at 2 cycles of 6,000 rpm for 20 sec and 2 cycles of 6,500 rpm for 30 sec with intermittent cooling between each cycle. Following centrifugation at 14,000 rpm for 10 min at 4°C, the supernatant was passed through a 0.22 µm nylon filter. Filtrate was evaporated to dryness using Speed Vac (Thermo Scientific, MA, USA). The dried extract was dissolved in 50 µl of sample buffer (6.5:2.5:1 Acetonitrile: Methanol: Water) and used for further analysis. Targeted analysis of SORA was performed using a multiple reaction monitoring (MRM) based approach. SORA parent ion (m/z 465.1) to daughter ion (252.1) transition was optimized and used for the MRM analysis. Briefly, samples (10 µl) were injected in the XBridge HILIC column coupled with a hybrid triple quadrupole/linear ion trap mass spectrometer (SCIEX 4000 QTRAP system) and Shimadzu Prominence binary HPLC pump (Shimadzu, Japan). Data were acquired on positive ionization mode. The samples were eluted at a flow rate of 700 µl/min with 32 min linear gradient starting from 5% mobile phase A (10 mM ammonium formate with 0.1% formic acid) and increasing to 60 % mobile phase B (acetonitrile with 0.1% formic acid). The column was kept at 60% mobile phase B for 3 min then returned to 5% mobile phase A for equilibration. MS conditions were set as follows, source temperature: 400°C, interface heater: on, curtain gas: 30, collision energy: 44, declustering potential: 90, entrance and exit potential: 10, and the two ion source gases were set at 45 arbitrary units. The standard curve was plotted with serial dilutions of pure SORA (ranging between 1 ng to 22 ng) versus the analyte peak area. Analyst 1.5 software (Sciex, CA, USA) was used for the analysis of acquired data. For the integration of peak areas, the Analyst quantitation wizard was used. The peak areas obtained after integration were exported to a spreadsheet file format.

### 2.10 Animal experiment

NOD/SCID, male mice (6-8 weeks) were procured from the in-house Experimental Animal Facility (EAF) at NCCS, Pune, India. After acclimatizing for one-week mice were divided into two groups. One group was fed on a normal diet (ND) and the other on a high cholesterol diet (HCD). HCD was prepared by adding 2% cholesterol dissolved in ethanol, followed by coating the solution on a normal diet which was further dried and autoclaved before feeding to mice. Water and food were always provided *ad libitum*. The mice were fed on respective diets for 2 months and these were injected subcutaneously with 5×10^6^ Huh-7 cells.

Thereafter, tumor formation was monitored by visualisation and the dimension of the palpable tumor was measured by using a digital vernier calliper at 3 days intervals. Tumor volume was calculated using the formula: 0.523 x (Length) x (Width)^2^. When the average tumor size reached 100 mm^3^, tumor-bearing mice were randomly distributed into experimental groups as per the protocol mentioned in figures 6 A and 7A, drug treatment was initiated.

SORA 20 mg/kg was administered orally using oral gavage and 7.5 mg/kg AVA was injected intraperitoneally (i.p.) into the mice. A fresh solution of SORA was prepared in solvent (12.5% ethanol; 12.5% kolliphor; 75% water), and AVA in DMSO under sterile conditions. Respective vehicle controls were administered to the control group. Serum was collected through retro-orbital puncture after giving anaesthesia (Xylazine: Ketamine,1:4). At the end of the experiment mice were sacrificed and tumors were excised. Samples were preserved at −80^о^C for further molecular studies. For histological studies, organs and tissues were preserved in 10% paraformaldehyde (PFA) at 4°C. The experimental layout is depicted in figure 6 A and figure 7 A. All the animal experiments were carried out as per the requirement and guidelines of the Committee for the Purpose of Control and Supervision of Experiments on Animals (CPCSEA), Government of India, and after obtaining permission from the Institutional Animal Ethics Committee (IAEC).

### 2.11 Experimental layout

When the tumor of all the mice reached an average volume of 100 mm^3^, mice were randomly distributed into four groups. One group (n=3 mice) was kept as drug vehicle control and the second group (n=3 mice) was administered with AVA (7.5 mg/kg), the third group (n=3 mice) was treated with SORA (20 mg/kg) and the fourth group (n=5 mice) was administered with the combination of both AVA (7.5 mg/kg) and SORA (20 mg/kg). Treatments were given daily until tumor size reached <2000 mm^3^.

### 2.12 Statistics

Values are represented as means ± SD. *, p < 0.05, **, p < 0.005, ***, p < 0.0005, ANOVA followed by Dunnett’s test, Student’s t-test where. *, p < 0.05, **, p < 0.01, ***, p < 0.001. Graphs were produced using GraphPad Prism 8 and Microsoft Excel.

## 3. Results

### 3.1 Hypercholesterolemia hampers the cytotoxic effect of SORA in HCC cells

To investigate the role of LDLc on the sensitivity of cells towards SORA, HCC cells were treated with SORA in the presence or absence of LDLc for 48 h and cell survival was assessed by MTT assay. LDLc exposure diminished the cytotoxicity of SORA in HCC cells (Figure 1A). Further, to explore whether a lower concentration of LDLc also has any effect on the sensitivity of HCC cells to SORA, Hep-G2 and Huh-7 cells were treated with SORA in the presence of 10 µg/ml LDLc and the long-term survival was compared with the cells cultured in the presence of 100 µg/ml LDLc. It was observed that higher LDLc concentration (hypercholesterolemia) abrogated the cytotoxicity induced by SORA as compared to the lower concentration of LDLc which remained comparable to the cell survival upon SORA treatment alone (Figure 1B and Supplementary Figure 1A). In addition to LDLc, another cholesterol carrier is high-density lipoprotein (HDLc). HDLc particles are involved in reverse cholesterol transport as they carry cholesterol from vascular tissues to the liver where eventually it is excreted in bile [21]. The presence of HDLc reduces the cytotoxicity of SORA and significantly increases the long-term survival of HCC cells as compared to the SORA treatment alone (Figure 1C and Supplementary Figure 1B). These observations suggested that extracellular cholesterol either in the form of LDLc or HDLc diminishes the cytotoxicity of SORA in HCC cells. To further understand the underlying mechanism; intracellular SORA accumulation was quantified by mass spectrometry analysis. It revealed that LDLc treatment in cells causes a significant reduction in the intracellular accumulation of SORA (Figure 1D). Overall, these observations indicate that hypercholesterolemia diminishes the cytotoxic potential of SORA which is partly a consequence of reduction in the intracellular accumulation of SORA.

**Figure 1.**
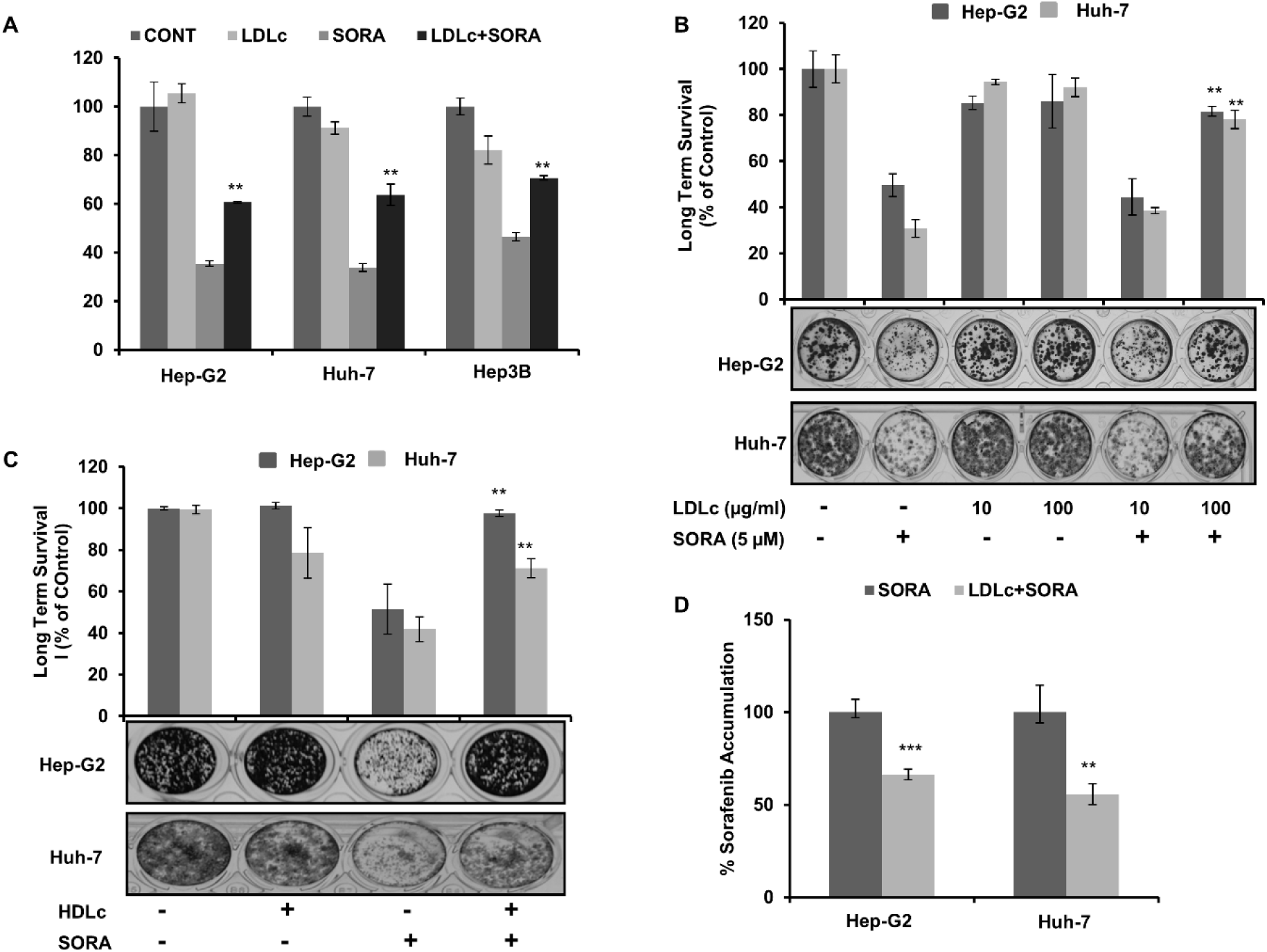
Hypercholesterolemia impairs the outcome of SORA treatment in HCC cells. (A) HCC cells (2 × 10^3^ cells) were plated and pre-incubated with LDLc (100 µg/ml) for 24 h followed by treatment with SORA ( 5 µM) for 48 h as indicated. Cell survival was assessed by MTT assay. (B) and (C) Hep-G2 and Huh-7 cells were pre-treated with indicated concentrations of LDLc and HDLc respectively for 24 h followed by SORA treatment for 48 h. Cell survival was assessed by long-term survival assay. (D) Intracellular SORA accumulation. Values are given as mean ±SD for SORA treatment vs. SORA + LDLc treatment. Experiments were done in triplicate and performed thrice. The values of *p < 0.05, **p < 0.01 and ***p< 0.001 were considered as statistically significant.

### 3.2. AVA enhances the cytotoxic ability of SORA under hypercholesterolemia

To investigate the combined effect of SORA and anti-hypercholesterolemic agents, the effect of AVA (ACAT inhibitor) and simvastatin (SIM) (hydroxy methylglutaryl CoA reductase inhibitor) on the cells was evaluated by MTT assay. It was observed that both drugs inhibited HCC cell survival in a dose-dependent manner (Supplementary figure 2A and B). Furthermore, to evaluate whether these cholesterol-lowering drugs have any consequence on the functionality of SORA, HCC cells were treated with AVA (5 µM) and SIM (5 µM) either alone or in combination with SORA (2.5 µM). The coefficient of drug interaction (CDI) was found to be less than 1 when cells were treated with the combination of SORA with AVA or SORA with SIM. The CDI value in SORA and AVA treatment group was observed to be less than the CDI value of SORA with SIM treatment (Supplementary table 1). Therefore, for all further studies, co-treatment of AVA with SORA was investigated.

Half inhibitory concentrations (IC_50_) of AVA were calculated in the absence or presence of LDLc. As evident LDLc impairs IC_50_ of AVA, in Hep-G2 and Huh-7 cells (Figure 2A). Based on these observations 5 µM or 10 µM AVA concentrations were used to investigate the combinatorial effect of SORA and AVA in the presence of LDLc. HCC cells were treated with SORA, AVA and their combination in the presence of LDLc for 48 h and long-term cell survival was analysed. As expected, in the combination group, cell survival was reduced to 20% and 40% in Hep-G2 and Huh-7 cells respectively as compared with individual treatments with SORA or AVA (Figure 2B). Upon combining the drugs in the presence of LDLc, synergistic reduction in cell survival was observed in Hep-G2 (CDI= 0.40) and Huh-7 (CDI= 0.63) cells. In contrast, treatment with SORA and SIM exhibited an antagonistic effect in Huh-7 cells while the synergistic effect was noted in Hep-G2 cells in the presence of LDLc (Supplementary table 1B).

**Figure 2.**
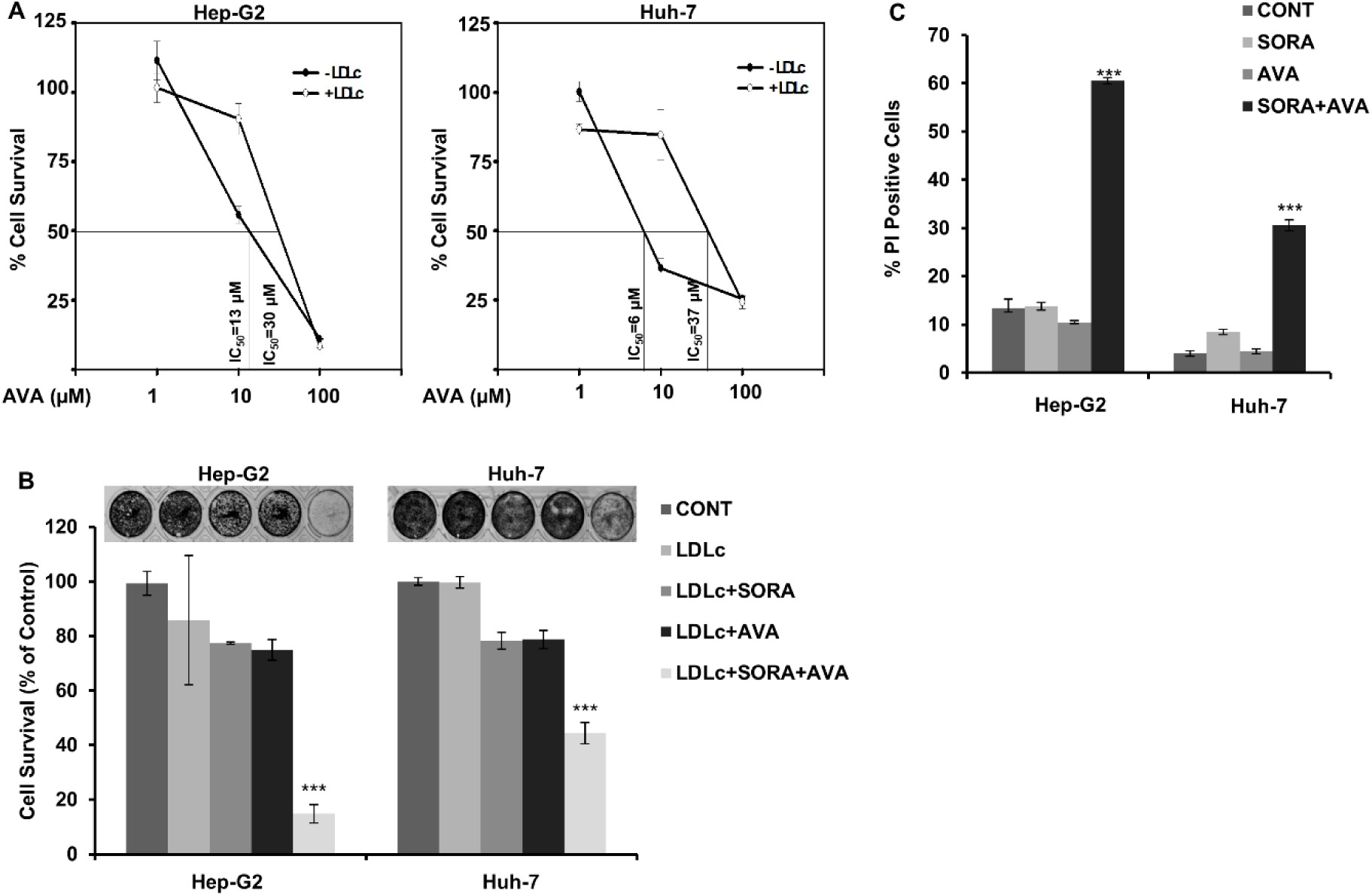
Combination of SORA and AVA shows synergistic effects. (A) MTT assay of Hep-G2 and Huh-7 cells (2 × 10^3^ cells) pre-incubated with LDLc (100 µg/ml) for 24 h followed by treatment with AVA for 48 h in the absence or presence of LDLc. (B) Long-term cell survival assay of Hep-G2 and Huh-7 cells pre-incubated with LDLc (100 µg/ml) for 24 h followed by treatment with SORA (5 µM), AVA (10 µM) and their combination for 48 h in presence of LDLc. (C) Quantification of dead cell population indicated by % PI-positive cells. Values are given as mean±SD. Experiments were done in triplicate and performed twice. The values of *p < 0.05, **p < 0.01 and ***p<0.001 were considered as statistically significant.

Further, the cytotoxic effect of SORA and AVA in the absence of LDLc was evaluated by staining the dead cell population with propidium iodide (PI) dye. In HCC cells increase in percentage of PI-stained cells was detected in drugs combination as compared to individual drug treatment indicating a synergistic cell cytotoxic effect (Figure 2C and Supplementry figure 3A). To validate the cytotoxic effect of SORA and AVA combination in the presence of LDLc, morphological changes in the cells were monitored. Microscopic examination revealed that HCC cells treated with the combination exhibited cell shrinking, rounding and detachment when compared with control (Supplementary figure 4).

**Figure 3.**
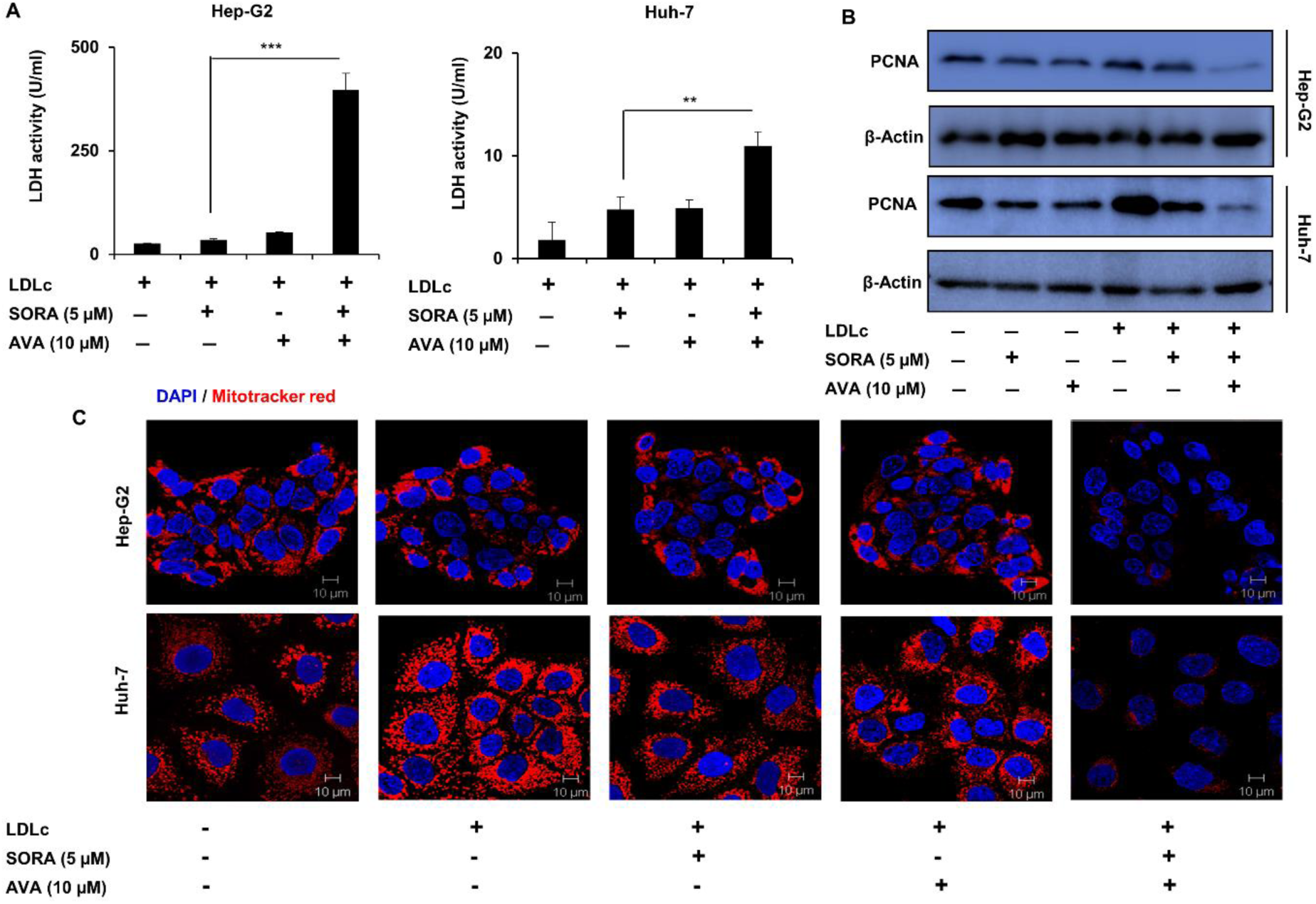
Cellular and molecular events associated with SORA and AVA-induced cytotoxicity under hypercholesteraemic conditions. (A) LDH release assay. (B) Hep-G2 and Huh-7 cells (3 × 10^5^ cells) were pre-incubated with LDLc (100 µg/ml) for 24 h followed by treatment with SORA and AVA for 48 h in presence or absence of LDLc for 48 h. Whole-cell lysates were subjected to Western blotting and the protein level of PCNA was analysed. β-Actin served as loading control. (C) Hep-G2 and Huh-7 cells were treated and processed for Mito Tracker Deep Red staining as described before. Bars represent 10 µm. Experiments were done in triplicate and performed twice. Blots were done once. The values of *p < 0.05, **p < 0.01 and ***p< 0.001 were considered as statistically significant. Confocal staining was performed twice.

Subsequently, the molecular events associated with anti-proliferative effects and cell death were evaluated by functional assays. The damage in the plasma membrane causes release of Lactate dehydrogenase (LDH) in the extracellular milieu and its activity is proportional to cell death [22]. In the presence of LDLc, the combination of SORA and AVA promotes the release of LDH as is evident by an increase in LDH activity in Hep-G2 and Huh7 cells (Figure 3A). Also decrease in the protein level of proliferative cell nuclear antigen (PCNA), was noted in cells treated with drugs combination as compared to individual drug treatments (Figure 3B). Earlier, it has been reported that alteration in mitochondrial outer membrane permeability and loss of mitochondrial membrane potential (ΔΨ) are lethal to cells and these are associated with cell death pathways [24,25]. It has been reported that active mitochondria uptake Mito-tracker Red dye in ΔΨ dependent manner and its fluorescence can be visualized under the confocal microscope. In the cells treated with SORA and AVA combination, a reduction in the fluorescence intensity was observed which is indicative of the loss of ΔΨ as compared to individual drug-treated cells (Figure 3C). These observations indicate that SORA together with AVA synergistically reduces HCC cell survival by promoting cell death because of an increase in plasma membrane permeability and loss of ΔΨ under hypercholesterolemic conditions.

### 3.3. Synergistic cytotoxicity of SORA and AVA combination was associated with downregulation of ERK activation and increase in ER stress

AVA is reported to inhibit ACAT isoforms (ACAT1 and ACAT2 which reduces the intracellular cholesterol esterification and storage of cholesteryl esters in form of lipid droplets (LD). By western blotting of cellular lysates ACAT1 and ACAT2 isoforms were detected in Hep-G2 and Huh-7 cells (Figure 4A). Previously, it has been reported that intracellular accumulation of LD increases in the cells that are exposed to an excess of fatty acids or lipoproteins [26,27]. AVA is reported to inhibit cancer cell proliferation and it induces cell death by abrogating intracellular CE accumulation [18]. It also inhibited LDLc uptake and lipid droplet (LD) formation in drug-resistant pancreatic cancer cells thereby promoting synergistic cell killing when combined with gemcitabine [19]. Therefore, in the present study, it was assumed that AVA-mediated reduction in LD could be one of the mechanisms associated with cell killing when combined with SORA in the presence of LDLc. To visualize LD accumulation inside the cells, the use of Nile red fluorescence dye has been documented earlier [28,29]. In the cells treated with LDLc expectedly, an increase in Nile red staining is observed suggestive of intracellular accumulation of LD compared to untreated cells. AVA exposure alone or in combination with SORA did not reduce Nile red fluorescence intensity as compared to LDLc-treated Hep-G2 and Huh-7 cells suggesting that LD accumulation remained unaltered upon combination drug treatment (Figure 4A). In contrary to our findings, previously it has been reported that when cells were pre-treated with AVA for 48 h followed by 3 h exposure to diI-LDL (10 µg/ml), a decrease in the intracellular red fluorescence was noted indicative of a reduction in diI-LDL uptake [18] (Supplementary figure 5). Putting together these observations, the present study suggests that AVA-led suppression of LD accumulation is unlikely to be involved in potentiating cytotoxicity of SORA in the presence of LDLc. This prompted us to investigate other possible molecular mechanisms by which AVA enhances the efficacy of SORA under hypercholesterolemic conditions.

**Figure 4.**
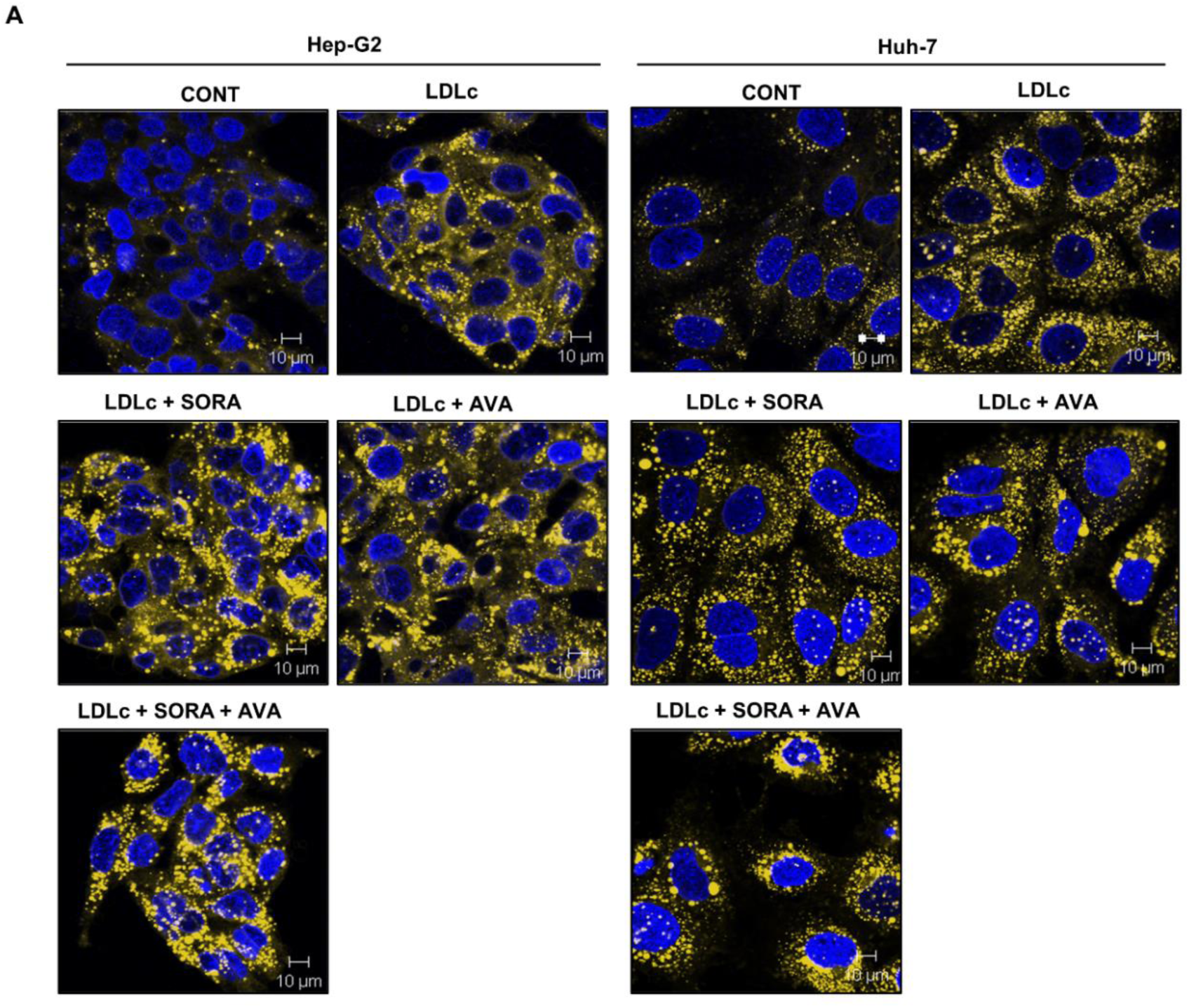
Combination of SORA and AVA does not reduce hypercholesterolemia-mediated LD accumulation. (A) Nile red staining to detect LD accumulation. Confocal stainings were performed twice. Blotting was performed once.

SORA is a multi-kinase inhibitor and suppresses RAF/RAS/MEK/ERK signalling. We have previously reported that hypercholesterolemia interferes with SORA-mediated p-ERK suppression [10]. To examine the effect of AVA co-treatment with SORA, the status of ERK activation was probed by western blotting. It was observed that ERK phosphorylation diminished in the cells treated with the drug combination as compared to individual drug treatments suggesting that inhibition of ERK activation could be associated with increase in cell death (Figure 5A-B). Further, to check whether AVA complements the p-ERK inhibitory effect of SORA and thereby, augments cytotoxicity, a potent and selective inhibitor of MEK/ERK kinase (U0126) was used in combination with AVA and cell survival was assessed [30,31]. AVA diminishes HCC cell survival when combined with U0126 in the presence of LDLc (Supplementary figure 6A-B). Studies have documented that SORA and AVA independently induce ER stress and promote cancer cell death. Therefore, we determined whether the combination drug treatment could induce ER stress in Hep-G2 and Huh-7 cells. The protein level of ER stress response marker; p-eIF2α was detected by western blotting after treating cells with either single agents or with the combination of SORA and AVA. Elevated p-eIF2α level in cells treated with drug combination is indicative of increased ER stress (Figure 5C-D). The protein level of transcription factor A mitochondrial (TFAM) which is involved in mitochondrial biogenesis did not change in HCC cells treated with either individual agents or drug combination s. This implies that the reduction in staining of Mito-tracker dye observed in figure 3C is unlikely to be due to a decrease in mitochondrial biogenesis and is likely to be a consequence of diminished ΔΨ. Collectively these findings indicate that cell death induced by AVA and SORA combination is likely because of reduced ERK activation and elevated ER stress in HCC cells.

**Figure 5.**
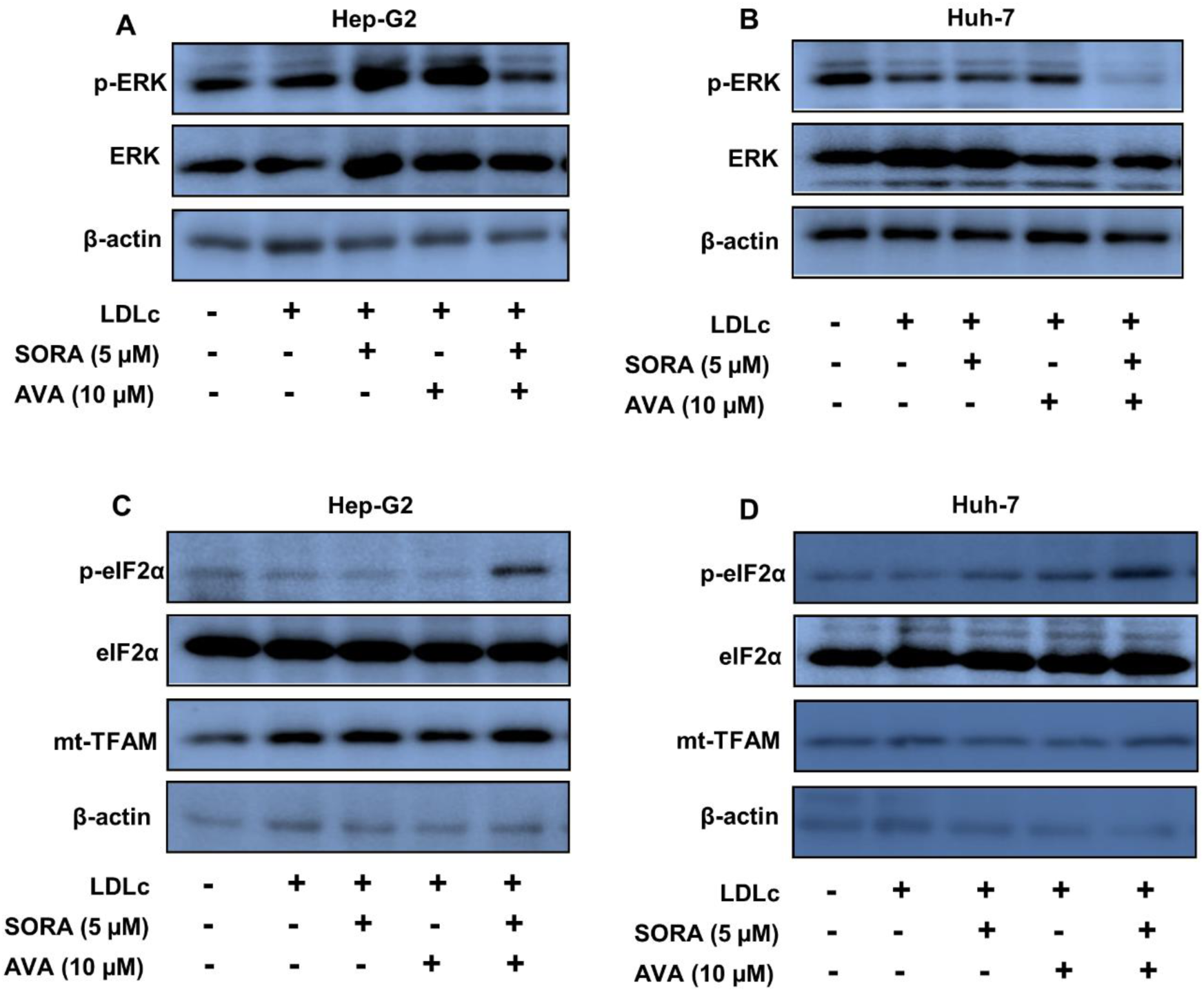
SORA and AVA combination diminishes ERK signalling and induces ER stress. (A) Western blot analysis for pERK, ERK, p-eIF2α, eIF2α and TFAM in Hep-G2 and Huh-7 cells (3 × 10^5^ cells) which were pre-exposed to LDLc (100 µg/ml) for 24 h followed by treatment with SORA (5 µM), AVA (10 µM) and their combination for 36 h. β-Actin was included as a loading control. Western blotting was done once.

### 3.4 The functionality of SORA in HCC is restored by the co-administration of AVA in HCD-fed mice

It has been reported that tumors grow rapidly in HCD-fed mice compared to ND-fed mice (33). Reduction in tumor weight and volume was detected in SORA-treated ND-fed mice compared to untreated mice (Figure 6A). To evaluate the consequences of hypercholesterolemia on the efficacy of SORA in HCC, mice were fed with HCD for 2 months followed by implantation of Huh-7 cells. Mice were grouped as described in the methods followed by treatment with either vehicle or SORA or AVA or with the combination of both (SORA + AVA), and the tumor progression was monitored. Individual treatment of AVA and SORA reduces tumor growth as compared to control. Interestingly, in the combination treatment group, SORA + AVA a drastic reduction in tumor growth was observed as compared to the single drug treatment group, which is evident by a decrease in tumor volume and weight (Figure 7C).

**Figure 6.**
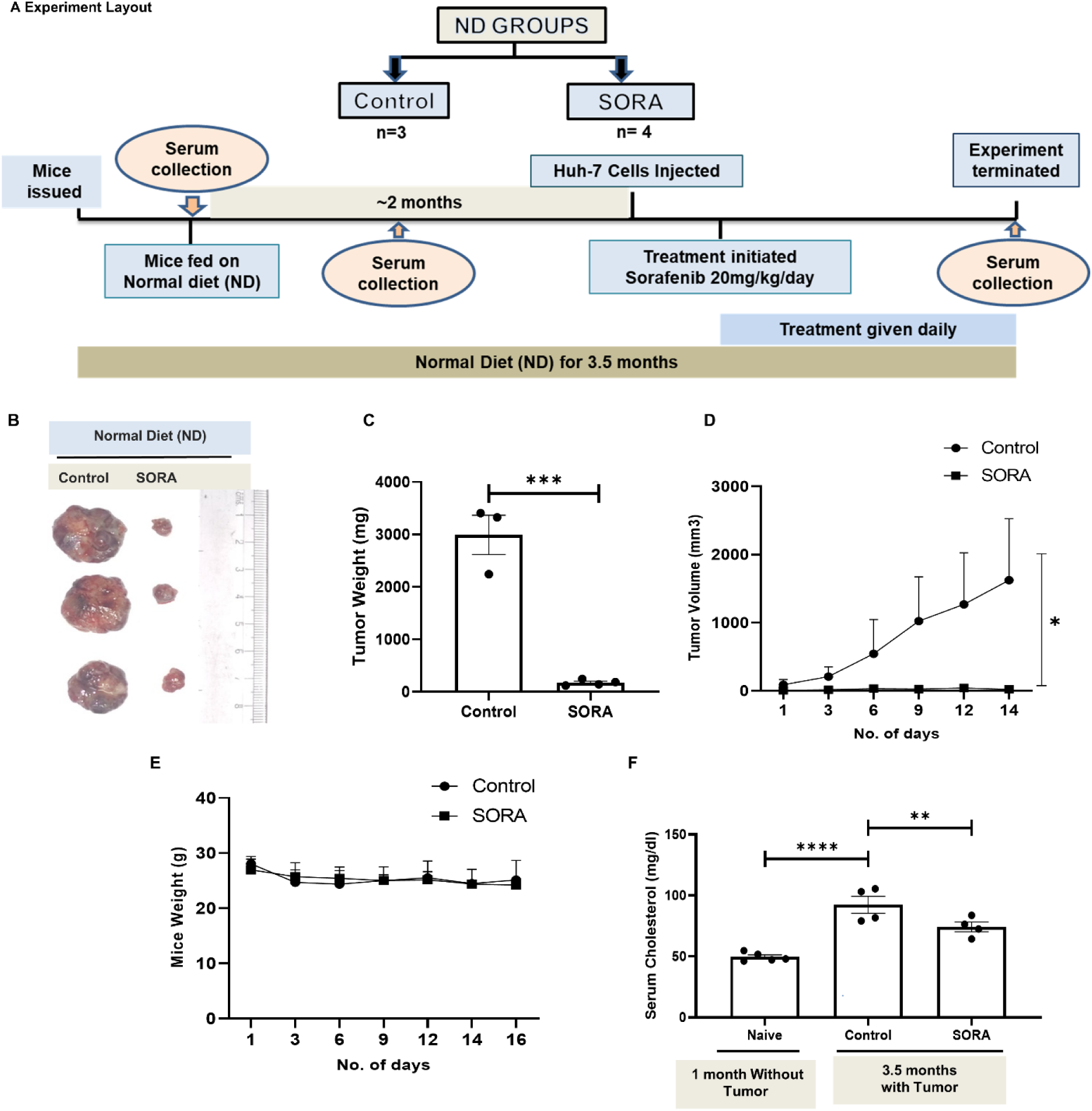
Depicts effect of SORA on tumor growth in mice fed on ND. A. shows experimental group layout and protocol timeline followed for Normal Diet (ND) mice. B. Tumor size pictures. C. Tumor weight. D. Tumor volume. E. Mice body weight. F. Cholesterol levels in serum. Experiment was performed once. The values of p < 0.05, p < 0.01 were considered as statistically significant (*, **) respectively

**Figure 7.**
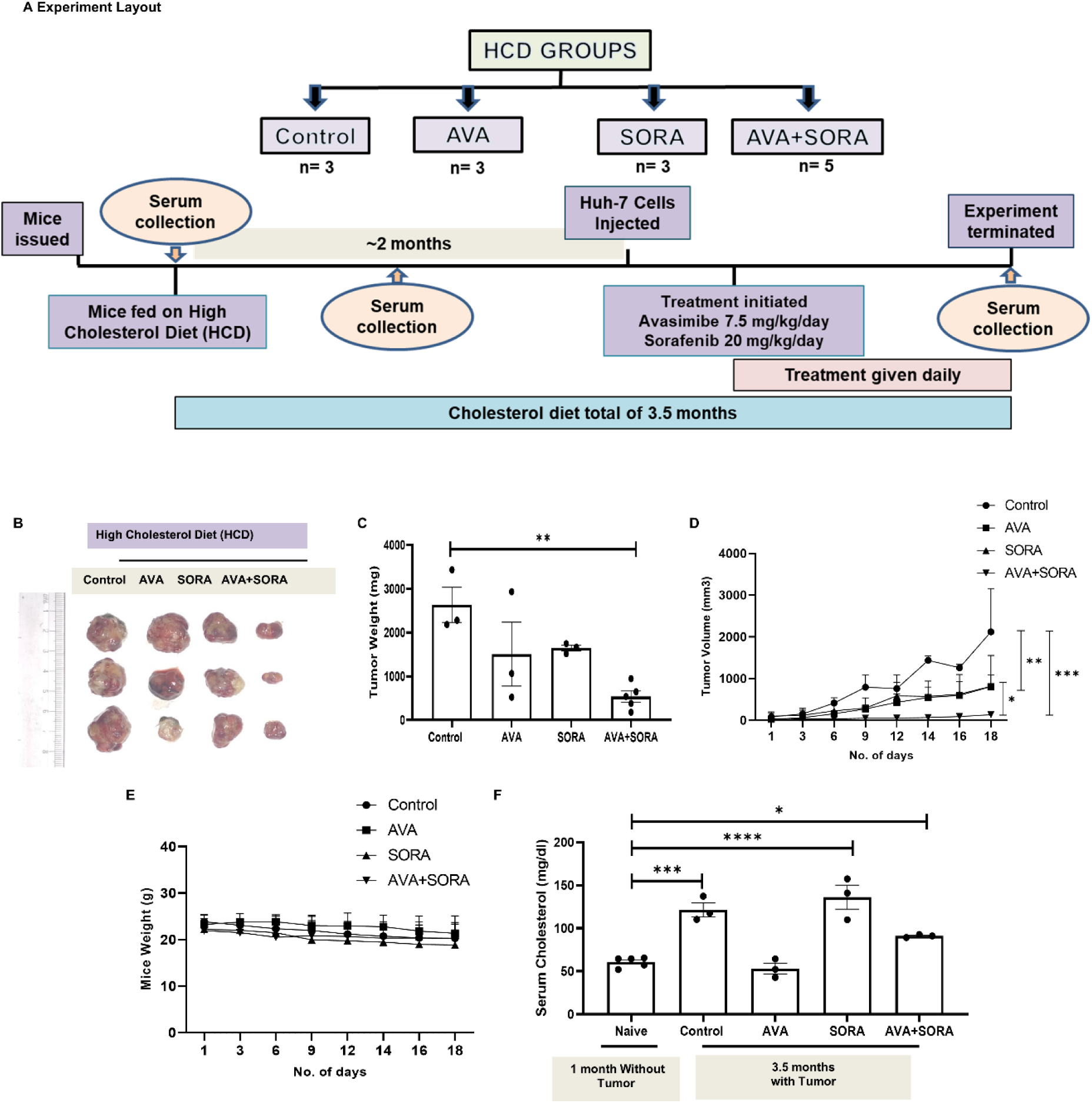
Depicts restored efficacy of SORA by co-treating mice with AVA in HCD fed mice. A. Experimental protocol timeline followed for High Cholesterol Diet (HCD) mice. B. Tumor size pictures. C. Tumor weight. D. Tumor volume. E. Mice body weight. F. Cholesterol levels in serum. Experiment was performed once. The values of p < 0.05, p < 0.01 were considered as statistically significant (*, **) respectively

**Figure 8.**
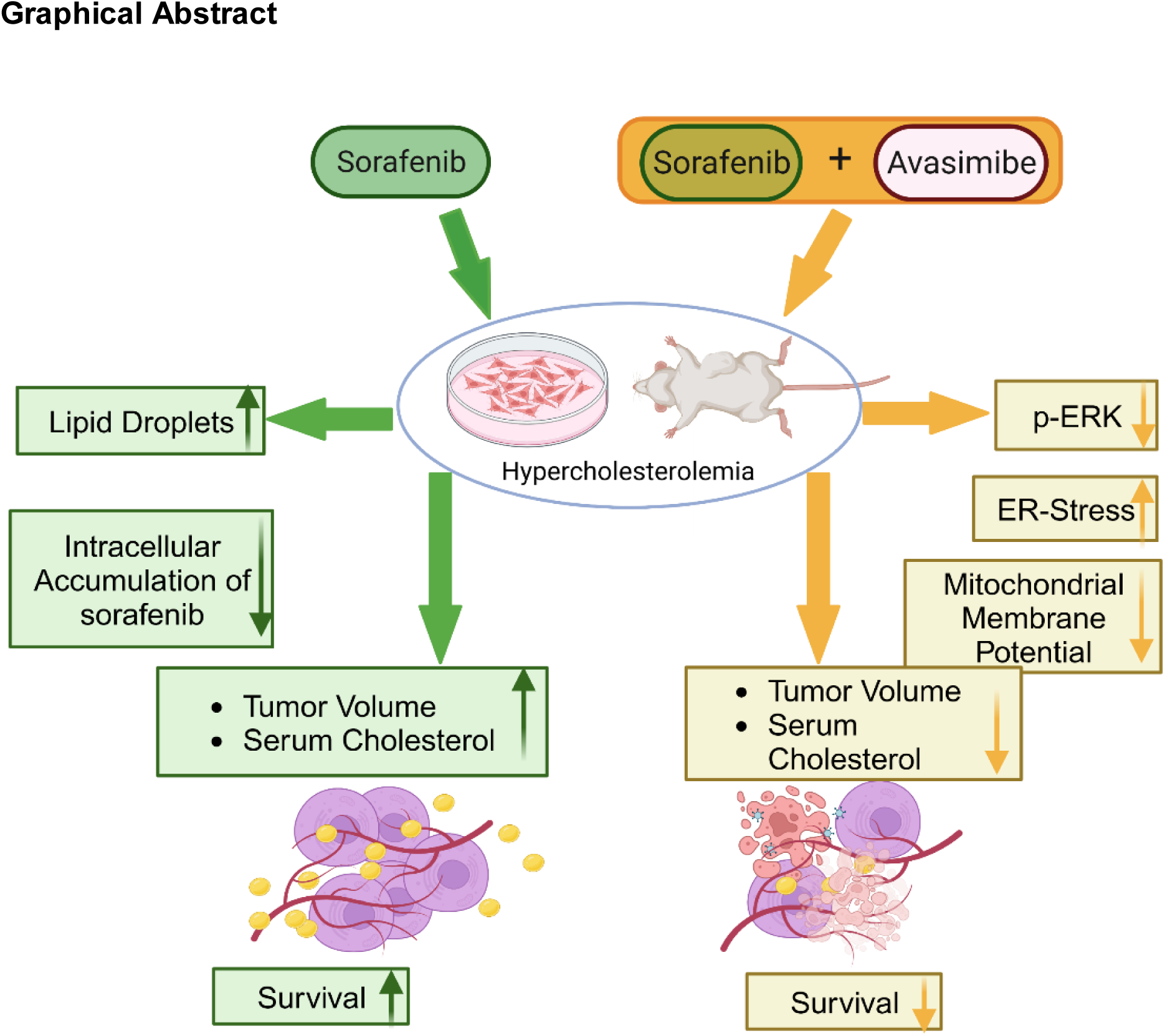
Schematic representing observations of the study. Hypercholesterolemia reduces the intracellular accumulation of SORA and promotes HCC cell survival. Co-treatment of HCC cells with SORA and AVA under hypercholesterolemia induces HCC cell death and is associated with reduced ERK signalling and elevated ER stress.

Cholesterol levels in serum were significantly enhanced in ND-fed tumor-bearing mice compared to mice without tumor. Also, cholesterol levels in serum decreased in tumor-bearing ND-fed mice treated with SORA compared to the control group (Figure 6). Interestingly, in AVA and AVA+SORA-treated group of mice, a significant reduction in serum cholesterol was observed. These results are indicative of AVA’s ability to control hypercholesterolemia induced by HCD (Figure 7F). No visible signs of toxicity were noticed in animals across all the groups as determined by measuring body weight during the course of the experiment (Figure 6). Therefore, it can be stated that under hypercholesterolemic environment AVA co treatment restors the functionality of SORA..

## 4. Discussion

Elevated circulatory cholesterol levels correlate with an increase in tumor growth, angiogenesis as well as metastasis in breast cancer and impairement in the immunosurveillance in pre-clinical colon cancer models [32–35]. Hwang et al., examined HCC patients (n=792) for cholesterol levels in serum and found 11.5% of patients (n=91) had hypercholesterolemia (252-973 mg/dl). The presence of larger tumors correlated with elevated cholesterol levels in serum [36]. In another similar study, it was reported that around 23% of total HCC patients had hypercholesterolemia (<220 mg/dl) [37]. Also, the cholesterol-rich extracellular milieu has been reported to alter the efficacy of chemotherapeutic drugs [8,22]. Previously, it has been shown that hypercholesterolemia has a negative impact on SORA efficacy. An earlier study from our lab indicated that LDLc abundance in the extracellular milieu reduces the intracellular accumulation of SORA in HCC cells [10]. It has been reported that SORA administered in combination with doxorubicin, 5-fluorouracil and other molecular targeted therapies did not lead to a promising outcome in HCC clinical trials [40]. Considering the dependency of cancer cells on lipogenic pathways, researchers have explored the implications of lipid-lowering agents like statins either alone or in combination with chemotherapy [41,42]. Statins potentiated SORA cytotoxicity in various cancer cell lines and in the preclinical model of HCC [23,43]. The current study unravels the efficacy of SORA in combination with anti-hypercholesterolemic drugs in the hypercholesterolemic HCC mouse model. In the present study, SORA combined with statin exhibited synergistic cytotoxicity. The efficacy was nullified in the presence of high LDLc levels. Recently, the anticancer property of AVA has also been explored either alone [54–55] or in combination with BCL-ABL-2 inhibitor imatinib [56]. To the best of our knowledge, this is the first study that demonstrates the synergistic cytotoxic effect of SORA and AVA combination on HCC in the presence of LDLc.

In HCD-fed mice, SORA treatment reduces tumor volume by half whereas the decrease is 2-fold higher in ND-fed mice compared to their vehicle controls. This indicates that increased serum cholesterol level in HCD-fed mice is likely contributing towards a decrease in the activity of SORA. Therefore, it can be stated that the administration of AVA along with SORA in HCD condition dramatically improves the efficacy of SORA.

The anticancer effect of AVA is dependent on ACAT [19,20,44]. A study on early-stage human HCC tissue samples suggested, ACAT as a targetable protein candidate. Also, AVA treatment reduced the growth of patient-derived tumor xenografts [44]. Therefore, in this study, we explored the involvement of ACAT in AVA-mediated restoration of SORA functionality. Previously, treatment with AVA has been shown to reduce LDL uptake in pancreatic cancer cells, and lipid content in 3T3-L1 preadipocytes during adipogenesis [18,46]. Hence, we hypothesized that AVA treatment could decrease hypercholesterolemia-led intracellular LD accumulation and improve the responsiveness of HCC cells to SORA. Surprisingly, AVA treatment as a single agent or in combination with SORA did not cause a reduction in LD content as detected by Nile red staining. As no change in LD accumulation upon AVA exposure was observed in our study, which is contrary to earlier reports [18], we believe these differences might be because of the sequence in which drug treatment was administered or due to the use of higher LDLc concentrations.

Activation of the RAS/RAF/MEK/ERK signalling cascade plays a role in HCC progression. Also, small molecule inhibitors such as Salirasib, Novartis, CI-1040, PD0325901, Selumetinib and AZD6244 etc., against this signalling pathway are beneficial in HCC management. [47,48]. Activated ERK (p-ERK) is a key effector molecule of this pathway and SORA treatment is reported to inhibit p-ERK in a dose-dependent manner [49,50]. Therefore, the involvement of the ERK pathway in mediating the cytotoxicity of the drug combination was investigated. Co-treatment of AVA with SORA resulted in diminishing p-ERK levels as compared to individual drug treatments. Additionally, when AVA was combined with another MEK/ERK inhibitor (U0126), HCC cell survival was decreased even in the presence of LDLc. This suggested that the downregulation of ERK activation participates in AVA-mediated increased responsiveness of HCC cells towards SORA upon LDLc availability.

Also, disturbances in lipid homeostasis can lead to ER stress [51]. It has been reported that in pancreatic cancer cells, AVA treatment induces endoplasmic reticulum stress and promotes apoptosis [18]. Similar to AVA, exposure to SORA is also reported to induce ER stress and apoptosis in HCC cells [52,53]. Therefore, the expression level of ER stress protein marker p-eIF2α was analysed in SORA and AVA-treated cells. though basal levels of eIF2α remain unchanged there was a significant increase in p-eIF2α protein level. Put together, increase in cytotoxic effect of combination treatment parallels with reduction in ERK phosphorylation and increase in ER stress.

In this study, in tumor-bearing mice increase in serum cholesterol level was observed compared to non-tumor-bearing mice which suggests that HCC progression contributes to alteration in serum cholesterol level and this finding is in agreement with the results obtained from clinical studies [36]. However, SORA treatment alone does not significantly influence serum cholesterol levels which may affect its efficacy. On the other hand, AVA causes a decrease in serum cholesterol levels and also attenuates tumor growth which suggests that AVA may function directly and indirectly in regulating the cholesterol metabolism of HCC tumors in hypercholesterolemic conditions. Impairment in the functionality of SORA towards controlling the growth of Huh-7 xenografted tumor in HCD-fed mice is indicative of an association between elevated cholesterol levels in serum with diminished effectiveness of SORA *in-vivo*.

## 5. Conclusion

The experimental results presented in this study together with available literature indicates that the complementation of AVA along with the primary drug SORA has the potential to reduce the growth of HCC under hypercholesterolemic condition. The growth inhibitory effect of this combination is achieved partly by regulating ERK activation and inducing ER stress.

## CRediT authorship contribution statement

**Dipti Athavale**: Conceptualization, Methodology, Data Curation, Formal Analysis, Investigation, Validation, Visualization, Writing - original draft, Writing - review & editing. **Himanshi Yaduvanshi and Firoz Khan Bhati**: Methodology, Investigation, Writing - original draft, Data Curation, Formal Analysis, Writing - review & editing. **Shyamananda Singh Mayengbam:** Investigation, Writing - review & editing. **Tushar More** Methodology, Investigation, **Srikanth Rapole** Resources, review & editing. **Manoj Kumar Bhat:** Supervision, Conceptualization, Funding Acquisition, Project Administration, Resources, Writing - review & editing.

## Abbreviations Used

HCC: Hepatocellular carcinoma
LDLc: Low-density lipoprotein cholesterol
HDLc: High-density lipoprotein cholesterol
SORA: Sorafenib
AVA: Avasimibe
SIM: Simvastatin
ND/HCD: Normal diet/High cholesterol diet
DMEM: Dulbecco’s Modified Eagle’s Medium
FBS: Fetal bovine serum
PBS: Phosphate buffer saline
PFA: Paraformaldehyde
DAPI: 4,6-diamidino-2-phenylindole
ANOVA: Analysis of variance
MTT: 3-(4,5-dimethylthiazol-2-yl)-2,5-diphenyltetrazolium bromide
ERK: Extracellular signal regulated kinase
ACAT: A cholesterol acyl transferase
LD: Lipid droplets
CDI: Coefficient of drug interaction
CE: Cholesteryl esters
PI: Propidium iodide
SDS PAGE: Sodium dodecyl-sulfate polyacrylamide gel electrophoresis
ER: Endoplasmic reticulum.
LDH: Lactate dehydrogenase
IC_50_: Half inhibitory concentrations
ΔΨ: Mitochondrial membrane potential
CPCSEA: Committee for the Purpose of Control and Supervision of Experiments on Animals
IAEC: Institutional Animal Ethics Committee
EAF: Experimental Animal Facility

