## Supplementry for "Hypercholesterolemia-induced impairment in sorafenib functionality is overcome by avasimibe co-treatment"

**Authors' affiliations:**

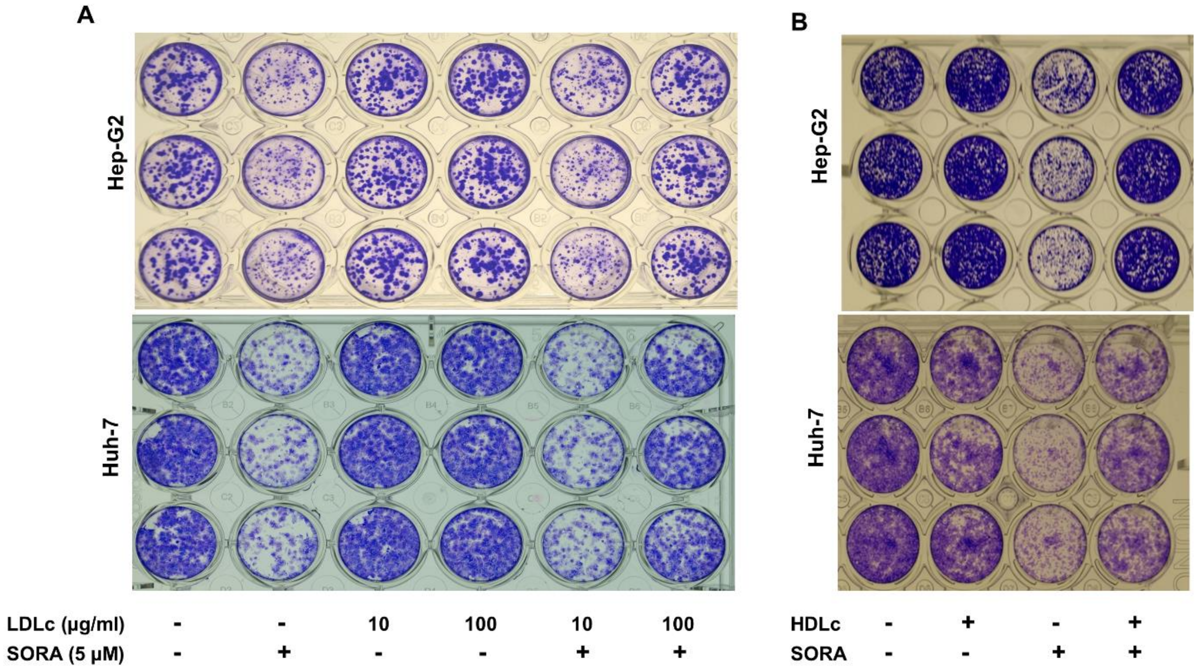


**Supplementary figure 1. Hypercholesterolemia impairs the outcome of SORA treatment in HCC cells (Huh-7 and Hep-G2).** (A) Long-term cell survival assay of Hep-G2 and Huh-7 cells pre-incubated with LDLc concentrations as mentioned for 24 h followed by treatment with SORA (5 µM) for 48 h. (B) Long-term cell survival assay of Hep-G2 and Huh-7 cells pre-incubated with HDLc (100 µg/ml ) for 24 h followed by treatment with SORA ( 5 µM) for 48 h. Experiments were done in triplicate and performed once.

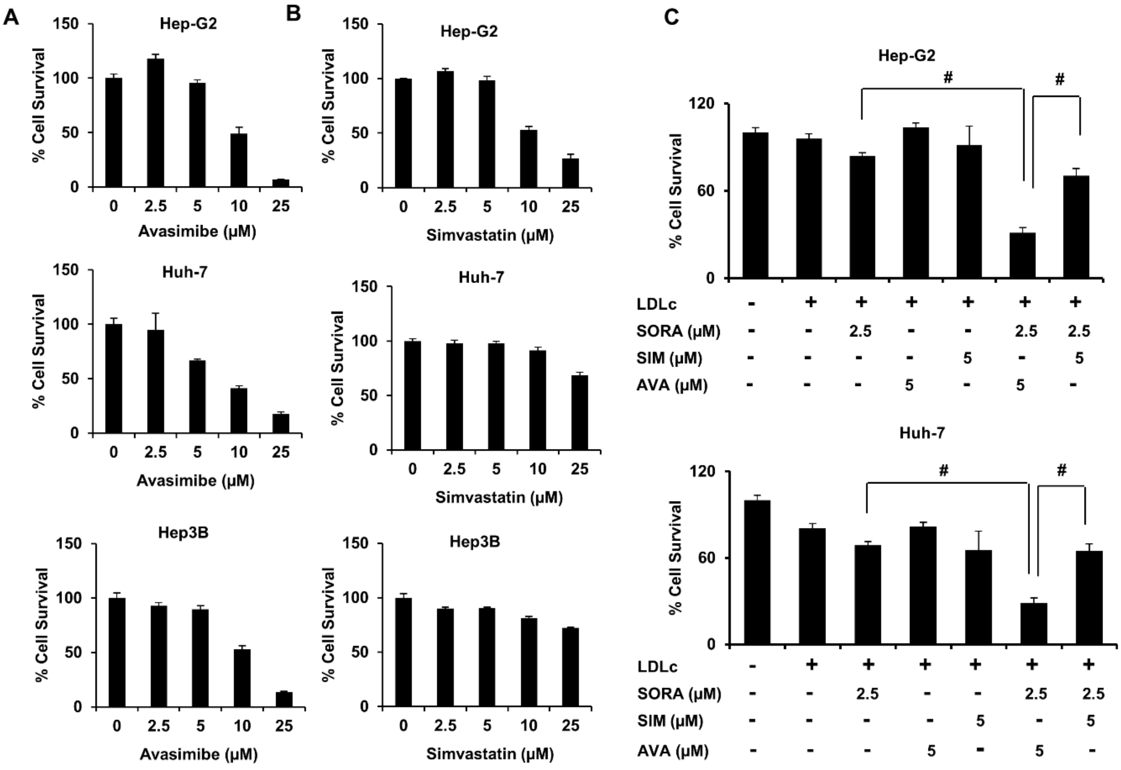


**Supplementary figure 2. Effect of varying concentrations of simvastatin and avasimibe on HCC cell survival.** (A) HCC cells (Hep-G2, Huh-7 and Hep3B) ($2 \times 10^3$ cells) were plated in 96-well plates and allowed to adhere. After 24 hours cells were treated with indicated concentrations of Avasimibe (μM) for 48 h, and cell survival was assessed by MTT assay. (B) Hep-G2, Huh-7 and Hep3B cells were treated with indicated concentrations of simvastatin (μM) for 48 h, and cell survival was assessed by MTT assay. (C) HCC cells ($2 \times 10^3$ cells) were plated in 96-well plates and allowed to adhere. Cells were pre-treated with LDLc (100 μg/ml) for 24 h in DMEM containing 5% FBS followed by exposure to the indicated concentrations of sorafenib (SORA), simvastatin (SIM) and avasimibe (AVA) for 48 h in presence of LDLc. Following treatment duration, cell survival was assessed by MTT assay. The results are given as means ± standard deviation of the representative experiment performed in triplicate. #p<0.0001 denotes significant differences. Experiments were done in triplicate and performed twice.

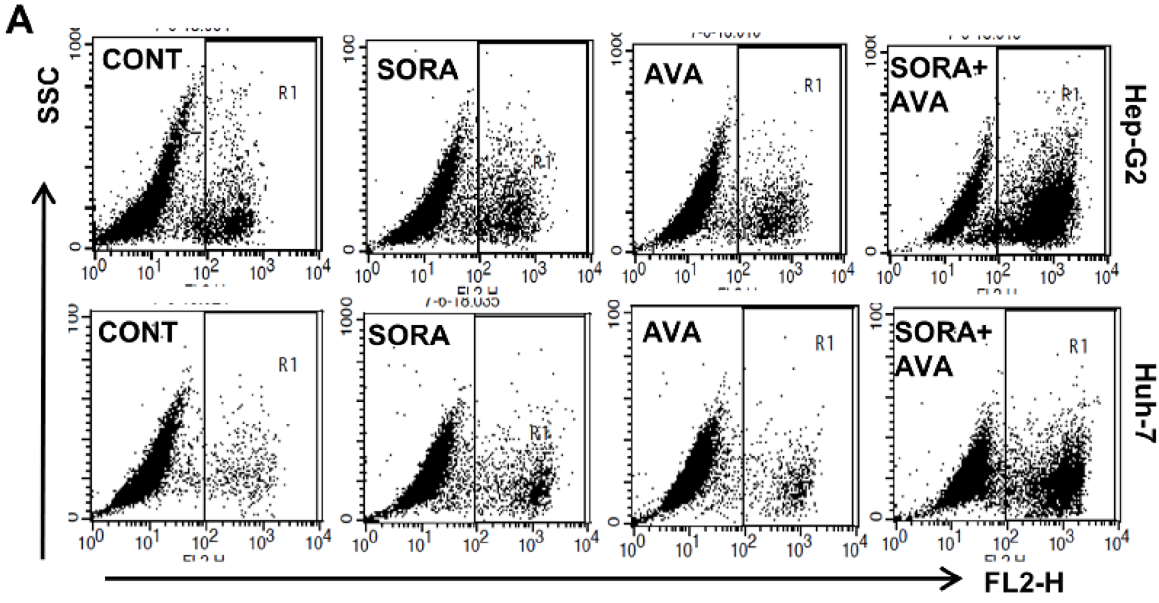


**Supplementary figure 3. Cytotoxicity of sorafenib and avasimibe combination on HCC cells.** (A) Hep-G2 and Huh-7 cells (3 × $10^5$ cells) were plated in 35 mm dishes and allowed to adhere. Cells were treated with sorafenib ( 5 μM) and avasimibe (10 μM) in DMEM with 5% FBS for 48 h. At the end of the treatment period, cells were trypsinized, washed with PBS, stained with PI and analyzed in FACS Calibur (BD Biosciences, USA). Scatter plots are indicative of one representative experiment. Experiments were performed twice.

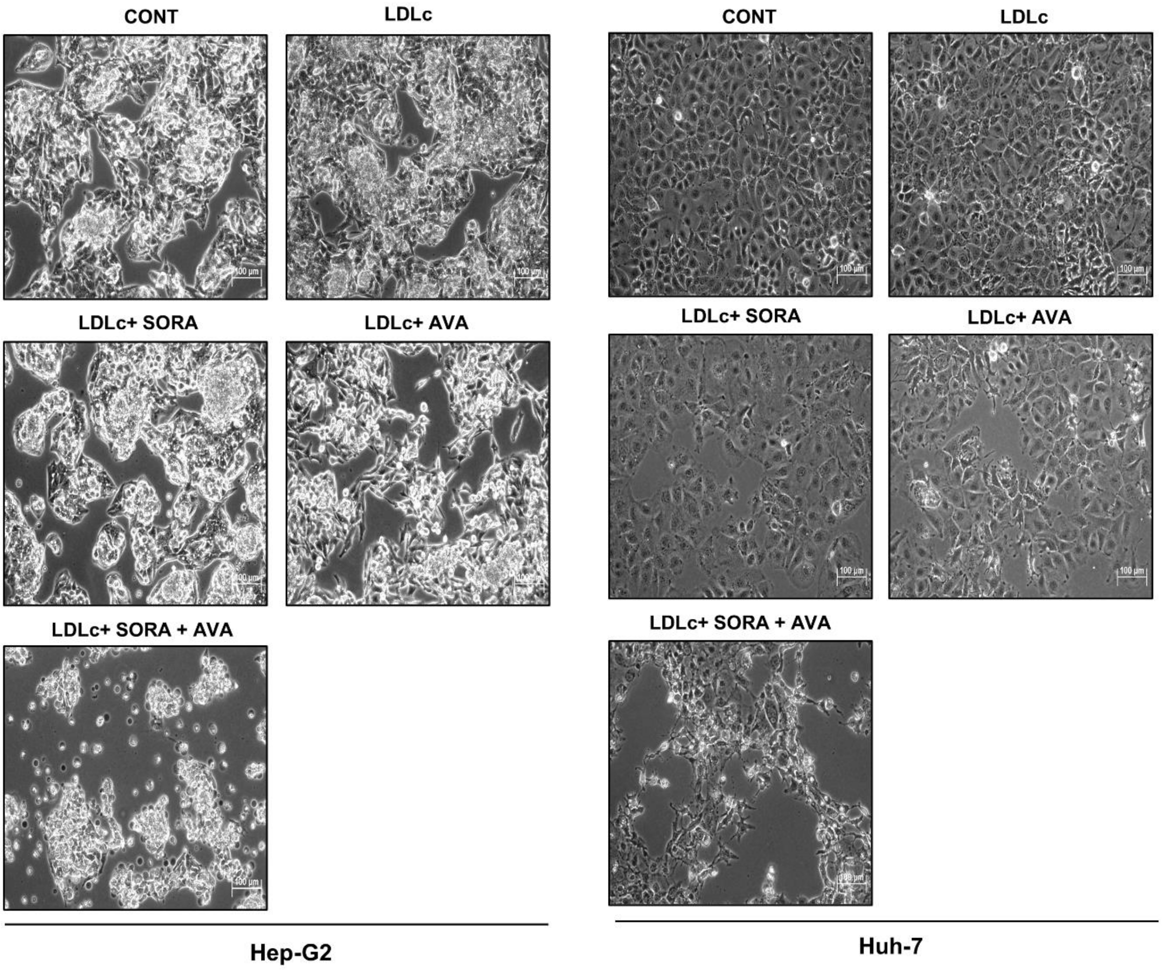


**Supplementary figure 4. Changes in cell morphology.** HCC cells (3 × $10^5$ cells) were plated in 35 mm dishes and allowed to adhere. Cells were pre-incubated in LDLc (100 µg/ml) for 24 h in DMEM with 5 % FBS followed by treatment of sorafenib ( 5 µM) and avasimibe (10 µM) for 48 h. At the end of the treatment period, cells were observed under a microscope and photographed with an attached camera at 20X. Bars represent 100 µm. Experiments were done in triplicate and performed twice.

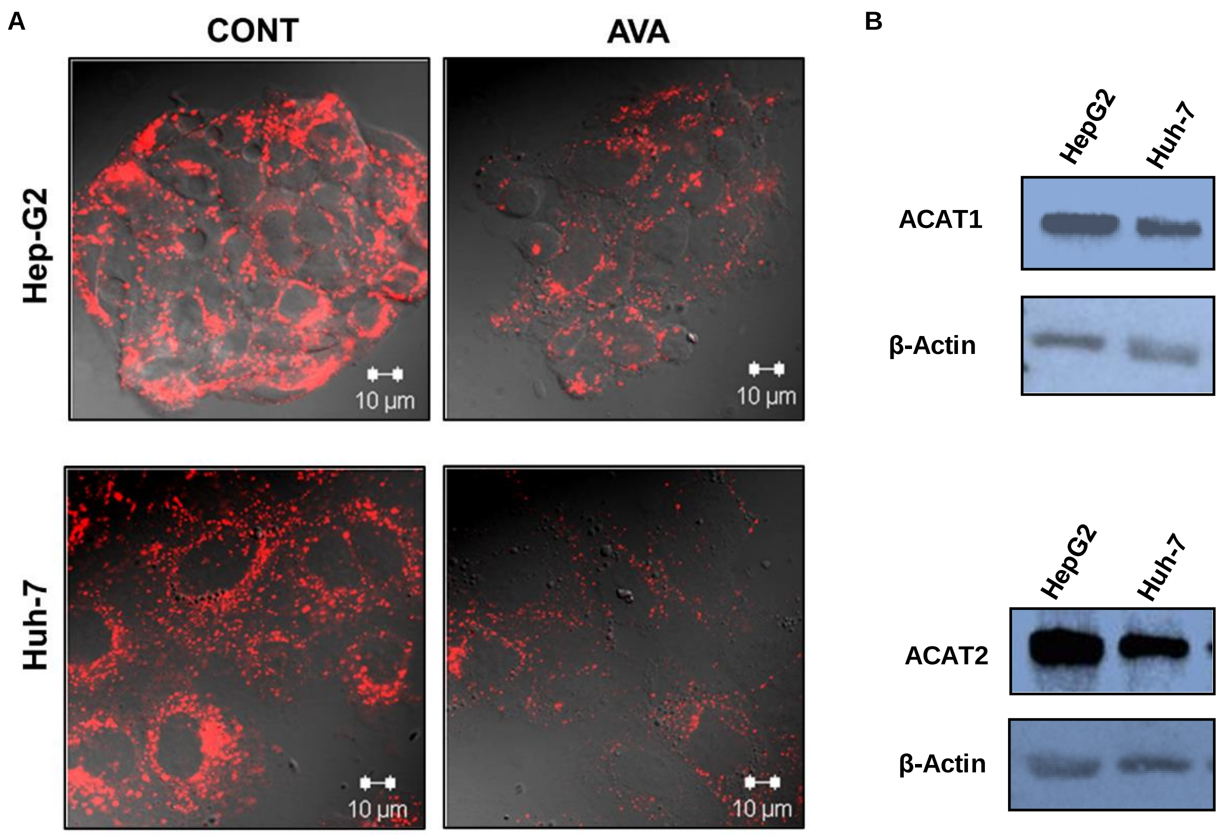


**Supplementary figure 5. Effect of avasimibe on DiI-LDL uptake.** (A) HCC cells were seeded on coverslips and treated with 10 µM avasimibe (AVA) for 48 h in DMEM with 10 % FBS followed by the addition of DiI-LDL for 3 h. Cells were washed with PBS and fixed with 4 % paraformaldehyde. Coverslips were mounted on slides and images were captured under a confocal microscope (Carl Zeiss, Germany). Bars represent 10 µm. Confocal staining was performed twice. (B) Western blot analysis for ACAT1 and ACAT2 expression in HepG2 and HuH-7 cells (3 X $10^5$) incubated in DMEM with 5% FBS for 48 h.

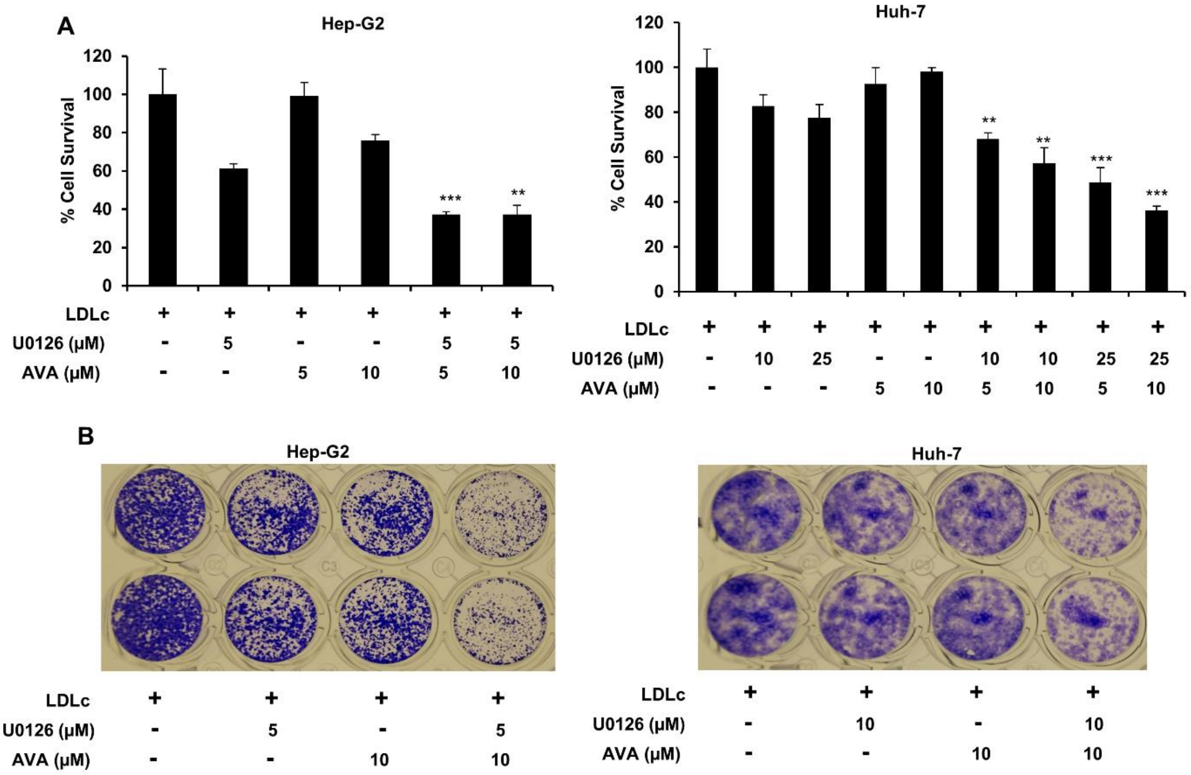


**Supplementary figure 6. Effect of MEK1/2 inhibition on the survival of HCC cells treated with AVA and LDLc.** (A) HCC cells (Hep-G2 and Huh-7) (2 × $10^3$ cells) were plated in 96-well plates and allowed to adhere. After 24 h cells were treated with indicated concentrations of LDLc, U0126 and Avasimibe (µM) for 48 h, and cell survival was assessed by MTT assay. The bar graphs are given as means ± standard deviation of the representative experiment performed in triplicate. The values of $*p < 0.05$, $**p < 0.01$ and $***p < 0.001$ were considered as statistically significant (B) HCC cells (Hep-G2 and Huh-7) were plated in 12-well plates and allowed to adhere. After 24 h cells were treated with indicated concentrations of LDLc, U0126 and Avasimibe (µM) for 48 h, and cell density was measured by crystal violate staining. Experiment is performed once in duplicate.

**A**

| | | Hep-G2 | Huh-7 | Hep3B |
|---|---|---|---|---|
| Growth inhibitory effect (OD) | Control (Vehicle control) | 1.7915±0.042 | 0.6183±0.024 | 1.0209±0.035 |
| | SORA (2.5 μM) | 1.1460±0.151 | 0.3746±0.006 | 0.8652±0.012 |
| | SIM (5 μM) | 1.4915±0.135 | 0.5834±0.021 | 0.8262±0.008 |
| | AVA (5 μM) | 1.5403±0.198 | 0.3764±0.016 | 0.6924±0.021 |
| | SORA + SIM | 0.9060±0.118 | 0.3393±0.010 | 0.6415±0.007 |
| | SORA + AVA | 0.5634±0.038 | 0.1961±0.012 | 0.3763±0.011 |
| CDI | SORA + SIM | 0.95 | 0.96 | 0.92 |
| | SORA + AVA | 0.57 | 0.86 | 0.64 |

**B**

| | | Hep-G2 | Huh-7 |
|---|---|---|---|
| CDI | LDLc+ SORA+SIM | 0.93 | 1.15 |
| | LDLc+ SORA+AVA | 0.40 | 0.63 |

**Supplementary table 1. Coefficient of drug interaction (CDI) OF SORA+SIM, SORA+AVA, LDLc+SORA+SIM and LDLc+SORA+ AVA.** (A) HCC cells (Hep-G2, Huh-7 and Hep3B) ($2 \times 10^3$ cells) were plated in 96-well plates and allowed to adhere. After 24 h, cells were exposed to the indicated concentrations of sorafenib (2.5 μM), simvastatin (5 μM) and avasimibe (5 μM) for 48 h. Growth inhibitory effect was assessed by MTT assay and CDI values are calculated (B) HCC cells (Hep-G2, Huh-7 and Hep3B) ($2 \times 10^3$ cells) were plated in 96-well plates and allowed to adhere. After 24 h, cells were pre-treated with LDLc (100 μg/ml) for 24 h followed by exposure to the indicated concentrations of sorafenib (2.5 μM), simvastatin (5 μM) and avasimibe (5 μM) for 48 h in presence of LDLc. The growth inhibitory effect was assessed by MTT assay and CDI values are calculated as mentioned before. CDI<1 indicates a synergistic effect, CDI=1 indicates an additive effect and CDI>1 indicates an antagonistic effect of a drug combination. The MTT assay was performed twice in triplicate**.**
